# No evidence of neural feature-specific pre-activation during the prediction of an upcoming stimulus

**DOI:** 10.1101/2025.09.16.676331

**Authors:** O. Abdoun, D. Todorov, A. Poublan-couzardot, C. Tirou, A. Lutz, M. Vernet, R. Quentin

**Author notes:** These authors contributed equally to this work (O. Abdoun, D. Todorov).

## Abstract

Our brains constantly make predictions about upcoming events based on prior knowledge of the environment. Although several neural mechanisms have been proposed to support this capacity, it is not yet clear how the brain makes such predictions. A compelling hypothesis is that the brain preactivates a sensory template of a predictable stimulus before it appears. In a recent study, Demarchi et al. had participants listen to sequences of sounds with different levels of predictability. For some sequences, participants could anticipate the next sound (in regular sequences), for others not (in random sequences). Using magnetoencephalography recordings and machine-learning methods to decode sounds from brain signals, Demarchi et al. concluded that auditory predictions pre-activate tone-specific neural templates before the sound onset. In our reanalysis of their data, we demonstrate that their results can be fully explained by a bias induced by the structure of the sequences: because the most likely stimulus also happens to be physically close to the previous one, spurious higher-than-chance decoding performance arises before the sound onset. We provide general criteria to assess whether a study is affected by this confound and requires a reexamination. We conclude that there is no evidence of anticipatory predictive perception in the Demarchi et al. dataset, and that existing evidence for feature-specific pre-activation during prediction in humans remains inconclusive.

## Introduction

It is now established that the brain predicts upcoming stimuli based on past experiences and that this predictive mechanism shapes perception^1^. The neural mechanisms leading to such predictions, however, are still unclear. One consensus is that such predictions are driven by top-down connections that move from higher-level regions to sensory regions. Predictions modulate neural signals in sensory regions before the predicted stimulus appears^1^. Based on these results, a potential candidate for neural prediction is the pre-activation, before the onset of a stimulus, of neural ensembles that encode the same stimulus during its perception^2,3^. To test this hypothesis, Demarchi et al.^3^ used magnetoencephalography (MEG) recordings in humans and multivariate pattern analysis (MVPA) to identify pre-activation of feature-specific neural activity during continuous and passive listening of four different tones in more or less predictable sequences (from random sequences to highly structured sequences). They trained MVPA classifiers to decode sounds from the MEG brain signals at the time of perception and tested these classifiers during the anticipatory period (i.e., MEG brain signals during the time preceding sounds’ onset). They found above chance level classification performance that correlated positively with sequence regularity, and concluded that the brain anticipates a predictable upcoming stimulus by pre-activating the neural population that encodes the same stimulus during perception.

In contrast, using the same open access dataset available^4^, we demonstrate through two different approaches that the reported classification performance is fully compatible with the absence of anticipatory feature-specific activity preceding sound onsets. First, we took MEG signals recorded during the passive listening of random sequences and re-ordered these signals to match the structured sequences (Figure 1a-b). Using these re-ordered signals that do not contain any possible anticipatory neural activity (because the sequence was random at the time of the recordings), we conducted the same analyses as in the original paper and identified the same effect that was interpreted as feature-specific predictive activity (Figure 1c). We then computed confusion matrices of the classifiers, which contain the probability distribution of the classifier outcomes conditioned on the true class. We found that classification errors were biased in favor of stimuli from neighboring frequencies. We further demonstrated that the misinterpreted predictive activity effect is caused by the combination of the transition probability matrices ruling the stimulus sequences and the biased structure of the classifiers’ confusion matrix (Figure 1e). Such an effect invalidates the 0.25 chance-level used as the null hypothesis by the authors. We conclude that there is no evidence of pre-activation of feature-specific neural activity during predictions in this dataset and briefly discuss the limitations of other existing evidence.

**Figure 1.**
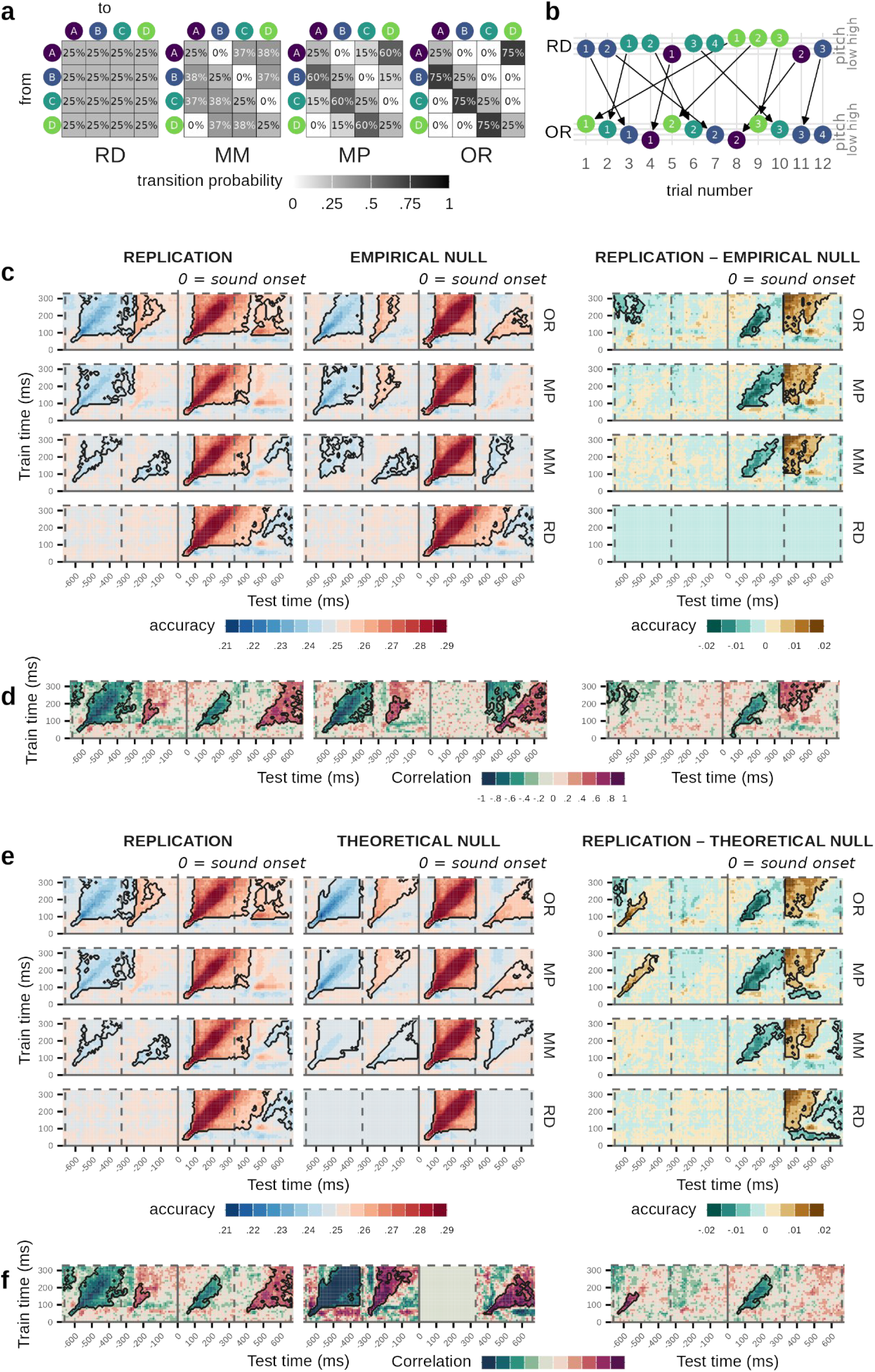
Time-generalization decoding of sounds presented in structured sequences with a classifier trained on random sound sequences. Statistically significant temporal clusters (p<.05) are delineated by a black line. **(a)** In the original study, four types of sound sequences were generated, each using a Markov chain characterized by a distinct transition matrix. The transition matrices were manipulated so that (i) probabilities of repetitions (diagonal elements) were set to 25% for all sounds and sequences, (ii) the identity of the most likely next sound given the current sound was the same in all sequences, and (iii) the entropy of the generated sequences decreased from RD (random) to MM (midminus), to MP (midplus) to OR (ordered). **(b)** In our reanalysis, we created a new control dataset by reordering data segments from random sequences (RD) according to the sequence order of lower entropy conditions (illustrated here with OR). By construction, any above chance level decoding accuracy on this control dataset could only be attributed to spurious sequence effects and not to predictive brain activity. **(c, Replication)** We replicated the approach of Demarchi et al. on the original dataset where above 25% chance level decoding performance appears several hundred of milliseconds before the stimulus (corresponding to the figure 3a in the original publication^3^). The display of our figure is different in two aspects from their original figure. First, we set up the color scale so as to reduce the saturation present in their original figure. Yet, in both the original and replicated analyses, the prestimulus decoding peaks in the highly regular condition (OR) at −230 ms (i.e., approximately 100 ms after the onset of the preceding sound) and exceeds the 25% chance level by less than 1 percentage point. Second, we have extended the testing time to cover the stimulus occurring two trials before the target stimulus, which revealed below 25% chance level accuracy, suggesting that decoding performance was at least partly driven by statistical dependency between stimuli. **(c, Empirical null)** is the same analysis, carried out on data extracted from the random (RD) condition and re-ordered to mimic structured sequences (see Methods and fig. 2b). Pre-stimulus decoding is found again, comparable in temporal pattern and magnitude to the original results as indicated by non-statistically significant performance in the contrast between the two analyses (right panel). Note that the positive upper diagonal during the subsequent stimulus (+333 ms to +666ms) is merely an artifact of the cropping used to create the reordered dataset. **(d)** Pre-stimulus decoding accuracy correlates positively with structure regularity, both in the replication of the original study and in the empirical null dataset. **(e)** Both the patterns of decoding performance and **(f)** correlations with statistical regularity are comparable to the replication results when the behavior of the classifier under the null hypothesis is calculated from the confusion and transition matrices (theoretical null).

## Results

### Reproduction of the original results

Demarchi et al.^3^ generated four types of sequences of varying entropy, from fully random to nondeterministic but highly-structured (Figure 1a). Following the methodology used in the original study (see Pipeline 1 in the method section), we trained decoding classifiers at the time of sound perception (from 0 to 333 ms after the sound onset) during random sequences (see supplementary figure 1 for the performance of these classifiers on random sequences). These classifiers were then tested on other time samples using the temporal generalization method^5^, including pre-stimulus interval, for every entropy level (Figure 1c).

A positive significant cluster of decoding performance is visible in the prestimulus interval (trial *n* ™ 1) for the most ordered sequence but not for the random one (Fig. 1c, Replication, −333 to 0 ms time-window). Moreover, the correlation between decoding performance and sequence predictability demonstrated a positive cluster during the prestimulus interval (Fig. 1d, Replication, same time-window). These results are similar to those reported in the original study, which were interpreted as the activation of a stimulus template before the stimulus appeared. During each sequence, 10% of the sounds were randomly replaced by omission trials with no sound at all. Results time-locked to these omissions also showed a significant cluster of decoding performance (above 0.25) during the pre-omission period in the ordered sequence (Supplementary figure 2a) and a significant cluster of positive correlation with entropy level during the same period (Supplementary figure 2b).

Increasing the temporal window of interest revealed negative significant clusters of decoding accuracy for the two most ordered sequences in the *n* ™ 2 trial interval, as well as negative correlation with predictability (Figure 1c & 1d, Replication, −666 to −333 ms time-window): a classifier trained to decode the sound at trial *n* have below-chance accuracy at the time of trial *n* ™ 2. This reflects the fact that the recorded brain activity at trial *n* ™ 2 is mostly evoked from the sound played at that time, and that the chance of having the same tone at trials *n* ™ 2 and *n* is low (i.e., 6.25%).

### Decoding accuracy under the null hypothesis

The authors of the original study have reached their conclusions by performing null hypothesis statistical testing (NHST) on the relationship between decoding accuracy and entropy level. NHST requires defining an operational null hypothesis, which is the expected value for the mean of a population parameter if the phenomenon of interest is absent. Here, the investigated phenomenon is the correlation *between sequence entropy, and the similarity between brain anticipatory activation patterns and sensory-evoked activation patterns*. The default null hypothesis for correlation coefficient is 0. Our reanalysis shows that this null hypothesis is incorrect in the context of the study. Our demonstration is based on two principled ways of generating accuracy data under the null hypothesis:

1. The *empirical null* approach tests the classifier on data epoched from random sequences and reordered so as to reproduce the sequences of stimuli actually presented to participants in structured sequences (figure 1b)
2. The *theoretical null* approach applies the confusion matrices of the linear decoder to the transition matrices of the structured sequences.

The two methods are conceptually equivalent, and both yield results that are statistically indistinguishable from the patterns reported by Demarchi et al., which they interpret as feature-specific prediction-related neural activity.

### Empirical null approach: Reproduction of the pre-activation patterns using re-ordered random sequences

In our replication of the results of Demarchi et al.^3^, the observed cluster of decoding performance during the interval of trial *n* ™ 1 in the ordered sequence (interpreted as anticipatory stimulus template activation in the original paper) seems to be time-locked to the preceding tone at −333ms. To test whether this cluster corresponds to stimulus template activation or is evoked by the preceding tone, we re-ordered the raw MEG signal acquired during the random sequence to match the three other entropy levels of the structured sequences (see Pipeline 2 in the method section). Using these re-ordered data (hence containing only brain response to unpredictable sounds), we used the same analysis as for the replication described above. Our results show a pattern of decoding performance similar to the original data (Figure 1c, Replication). Positive significant clusters of decoding performance are visible in the prestimulus interval (trial *n* ™ 1, −333 to 0 ms) for the two most ordered sequences and negative significant clusters in the preceding interval (trial *n* ™ 2, −666 to −333 ms) are visible for the three most ordered sequences (Figure 1c, Empirical null). The correlation across entropies of decoding performances for all time points of the time-generalization matrices demonstrated the same positive cluster during the prestimulus interval as in the original analysis (Figure 1d). The contrast between the original results and those obtained using the reordered random data failed to show a significant positive cluster during the prestimulus interval (Figures 1c and 1d, Replication – Empirical null). Furthermore, Bayesian analysis provides moderate evidence against a difference between the empirical null and replication patterns of correlations (Supplementary Figure 3), demonstrating that the positive decoding performance during the prestimulus interval in Demarchi et al. cannot be due to a neural prediction of the upcoming stimulus. We observed a positive upper diagonal during the processing of the subsequent stimulus (+333 ms to +666 ms) (Fig. 1d, contrast). This effect is unrelated to neural prediction and arises because the stimulus presented at 0 ms continues to be processed beyond 333 ms, but this ongoing activity is cropped when epochs are reordered.

Similarly, results time-locked to omissions in the reordered random data showed significant clusters of decoding performance (above decoding performance of 0.25) during the pre-omission period in the two most ordered sequences (Supplementary figure 2a, Replication) and a significant cluster of positive correlation with entropies during the same period (Supplementary figure 2b, Empirical null). Again, the contrast between the original results from omission trials and those deriving from the reordered random data failed to show a significant positive cluster during the prestimulus interval (Supplementary figure 2a, Replication – Empirical null), further demonstrating the absence of feature-specific prestimulus activation.

### Theoretical null approach: Reproduction of the pre-activation patterns using theoretical accuracy under the null hypothesis

During the prestimulus interval (trial *n* ™ 1), the chance of having the same tone as that in trial *n* (two identical consecutive tones) is 1/4; thus, theoretically, the classifier trained during random sequence at trial *n* should produce a chance-level accuracy of 0.25 when generalized during the interval of the trial *n* ™ 1 in the absence of feature-specific predictive neural activity. However, this reasoning holds true only if the confusion matrices of the classifiers are unbiased toward specific tones when making errors.

To test this assumption, we extracted the confusion matrices *C* from within-subject trained classifiers for each cell in the time-generalization matrix during the stimulus period, and calculated the probability *P* of correctly identifying stimulus *n* from stimulus *n* ™ 1 by chance only (see Pipeline 3 in the Method section):

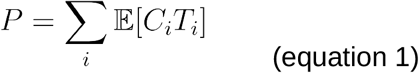

where *T* is the transition matrix of a given ordered sequence, *T*_*i*_ the *i*-th row of *T*, and *E* denotes mathematical expected value.

To understand how this true, theoretical chance level, deviates from the value of 0.25 assumed as chance level by Demarchi et al., consider the following rewriting of the covariance between the transition and confusion matrices:

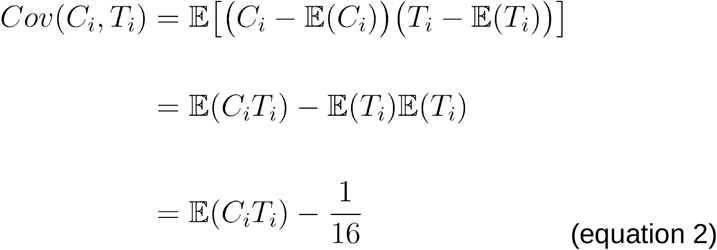

Combining it with the expression of the chance level above, we find:

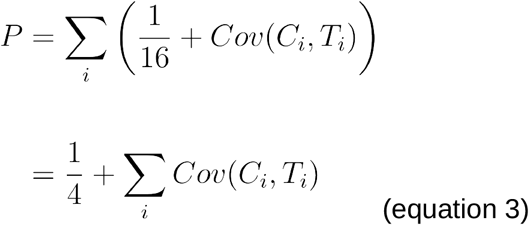

The actual chance level deviates from the presupposed value (one over class number, i.e. ¼ here) by the summed covariances between the rows of the transition and confusion matrices. In the sequences chosen by Demarchi and colleagues, the transition and confusion matrices are correlated because (i) sounds are more likely to be followed by the next lower pitch than by more dissimilar sounds (Figure 1a) and (ii) classifiers have more chance to misclassify two physically close sounds (Supplementary Figure 4).

We reproduced the same analysis as presented in Figure 1c and 1d using only these theoretical chance-level results (figures 1e and 1f). As expected, this method gave the same results than the reordering method of the random data (see pipeline 2 in Methods), producing clusters of decoding performance significantly above 0.25 during the prestimulus interval (figure 1e, Theoretical null) that correlated positively with sequence regularity (figure 1f, Theoretical null). These results are statistically indistinguishable from the patterns reported by Demarchi et al.^3^ that they interpret as feature-specific prediction-related neural activity (figure 1e, right and 1f, Replication – Theoretical null). Furthermore, Bayesian analysis provides moderate evidence against a difference between the theoretical null and original patterns of correlations, especially in the putative preactivation temporal cluster (Supplementary Figure 3).

### Decoding the most likely stimulus instead of the presented one

In the original study, the authors aimed to decode the presented stimuli before its onset. In non-random sequences, the presented stimulus was of course more often than not the most likely stimulus given prior information provided by the previous sound. For example, in midplus sequences (MP), the presented stimulus had a 60% chance of being the most likely, given the preceding sound. It might be argued that decoding the identity of the actual stimulus rather than the most likely one is somewhat unfair to the brain’s predictive machinery. Indeed, while the strength of neural predictions might be modulated by contextual predictability, the brain can not determine whether the next stimulus will obey the regularity or violate it ahead of the event. Following this reasoning, a more accurate assessment of the presence of neural anticipatory activation in response to structured sequences would be the decoding of the *most likely*, rather than *actual*, identity of the next stimulus. Thus, the approach used by Demarchi et al.^3^ might have attenuated the evidence for feature-specific, perceptual-like preactivation. We tested this possibility by replicating the same analysis as in figure 1c, but aiming to decode the most likely stimulus presented at *t*=0 given the identity of the previous one and the transition probability matrix ruling the sequence structure (figure 2a). After controlling properly for statistical dependencies using reordered random data (see Methods and figure 1b), we found a residual decoding accuracy above chance level which does not withstand cluster-based permutation tests (figure 2b, diff). Such non-significant decoding was of comparable absolute magnitude than the spurious effect found in the original paper (about 1% percentage point above chance level; figure 2b).

**Figure 2.**
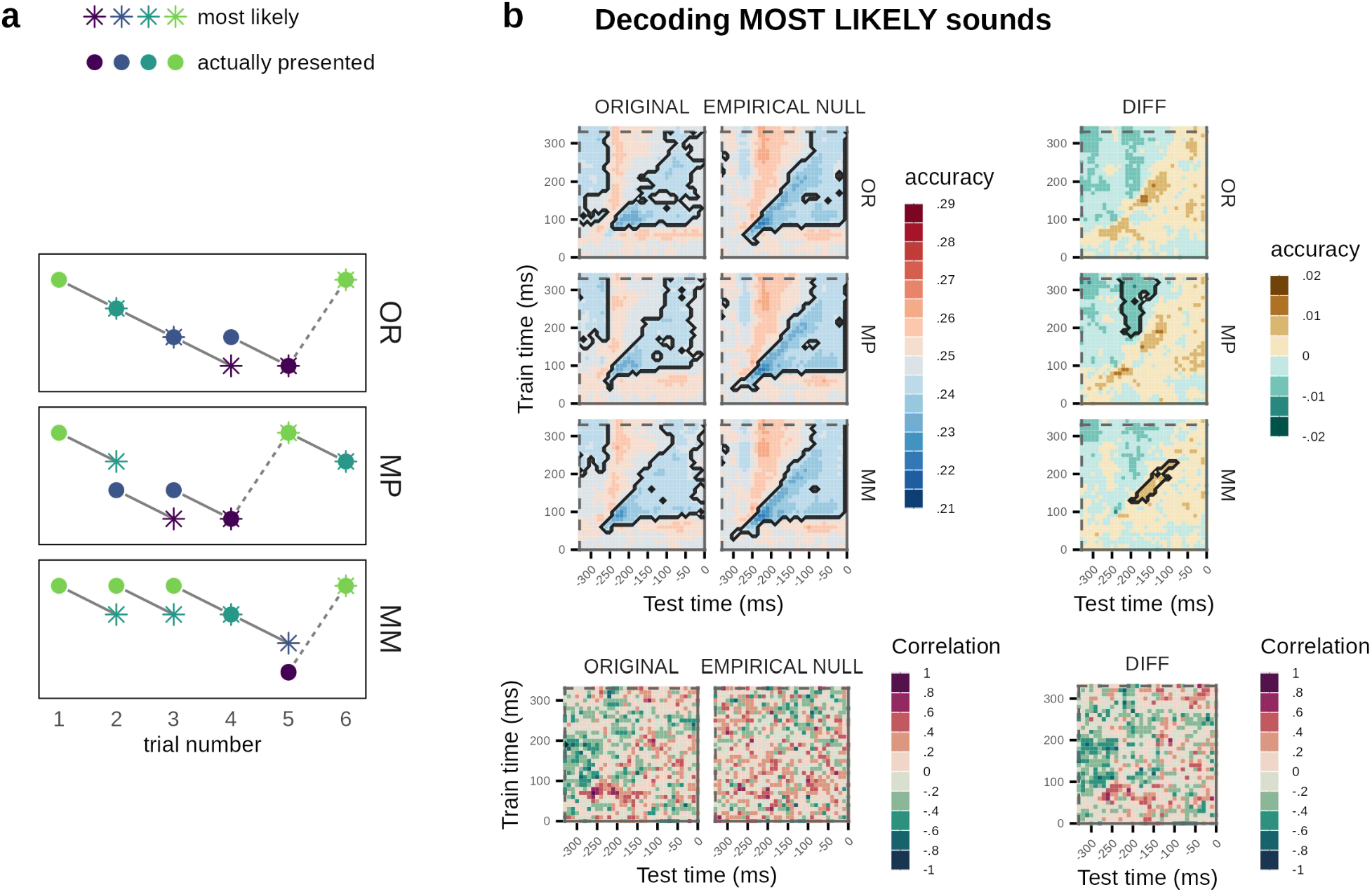
Time-generalization decoding of the sound most likely to be presented at t=0, given the identity of the previous sound, with a classifier trained on random sequence data. Compared to the original approach (replicated in figure 1c, left panel), this approach assumes the alternative and more plausible hypothesis that the brain holds the strongest predictive representation for the most likely stimulus, which is not always the actually presented one (see Methods). **(a)** Example of mismatch between the most likely sequence and the presented sequence. Demarchi et al. aimed at decoding the sound that was actually presented to the participants (colored circles), which was not always the most likely one given the previous stimulus (open gray circles); the lower the entropy, the more discrepancy between the two (MM > MP > OR). **(b)** When decoding the most likely stimulus, direct comparison of performance with the 25% chance level appears to suggest negative performance, but this is largely due to statistical dependency between the most likely and the actual sound, as is confirmed by decoding reordered data extracted from random sequences. After proper control of statistical dependency effects, only a weak (<1% point), non-significant decoding accuracy above the 25% chance level remains before stimulus onset. This effect does not correlate significantly with sequence entropy.

## Discussion

We demonstrated that there is no evidence of a feature-specific prediction-related neural activity in the human auditory system based on the open access dataset of Demarchi et al.^4^ (Figure 1). We also tested an alternative hypothesis in which the brain predicts only the most probable upcoming stimulus but this also failed to evidence a stimulus template activation during the prediction (Figure 2). Demarchi et al.^3^ have reached their conclusions by performing null hypothesis statistical testing on decoding accuracy, setting the chance level at 0.25 (i.e., one over the number of classes/possible tones). However, the assumed value of the null hypothesis was incorrect. The true null hypothesis is actually higher than 0.25. We demonstrate it by reconstructing a true null hypothesis in two ways: (i) by training our linear classifier the same way as in Demarchi et al.^3^, but testing it on data coming from random sequences and reordered stimulus-wise so as to reproduce the sequences of stimuli actually presented to participants in low entropy conditions (Pipeline 2 in the method section) and (ii) by extracting the confusion matrices from the same linear classifiers, and applying it to the sequences of stimuli presented to participants in low entropy conditions in a data-independent manner (Pipeline 3 in the method section). Both methods yielded similar results and invalidated the interpretation of a prestimulus activation of the stimulus neural pattern.

In the ordered sequence, having sound #1 at trial *n* ™ 1 means that the sound at trial *n* will be sound #4 with a 3/4 chance or sound #1 with a 1/4 chance. In the original analysis of Demarchi et al.^3^, their goal was to decode a stimulus before its onset with linear classifiers trained during the perception of a random sequence of sounds. This analysis makes an underlying assumption that the brain creates an internal model of the transition matrix and activates during the prestimulus interval a mixture of the stimulus neural template of stimulus #4 and #1. A simpler possibility is that the brain only pre-activates the most probable stimulus, in that case stimulus #4. We tested this hypothesis by attempting to decode, during the prestimulus interval, not the actual stimulus at time 0 ms, but the most probable one based on trial *n* ™ 1. While the prestimulus decoding performances were closer to significance when doing the differences between original and reordered random data, we did not find any evidence of a prestimulus feature-specific neural activity within this simpler hypothesis.

Kok et al.^2^ also proposed that predictions were associated with the generation of a stimulus template in sensory cortices before the onset of the stimulus. The presented visual stimulus (one out of two possibilities) was predicted with a 75% probability by a preceding sound (valid condition). In 25% of the cases, the other visual stimulus was presented (invalid condition). Linear classifiers were trained beforehand on random presentations of the visual stimuli, and then tested on task data, before the onset of the predictable visual stimuli. Kok et al. observed what they interpreted as a feature-specific neural activity only when contrasting valid *vs*. invalid trials. Crucially, the above-chance (respectively, below-chance) prediction-related activity was absent from the valid condition alone (respectively, the invalid condition alone). Moreover, such feature-specific neural activity began just 40ms before the visual stimulus onset and persisted for several seconds afterward, i.e., when the stimulus was clearly visible. Taken together, these characteristics of the reported effect cast doubt on whether this effect is specifically related to prediction mechanisms.

To summarize, the results of our reanalysis of the Demarchi et al.^3^ dataset concur with the observation that previous evidence of feature-specific neural activity^2^ is weak. Implications of our findings on recent studies making use of the Demarchi et al. paradigm will need to be carefully pondered^6–8^. More generally, the analytic expression for performance bias in equation 3 was derived under very few assumptions, making it applicable to a broad range of experiments: any experiment featuring a finite set of equiprobable stimuli presented through a first-order Markov process and analyzed using classifiers trained on neural responses to random stimuli. Notably, this bias holds regardless of sensory modality or classification method. Thus, studies within this scope should be reassessed and corrected for potential bias. On the other hand, studies using classifiers trained on data containing non-perceptual information (e.g., Schubert et al., 2023) cannot conclusively support interpretations involving sensory template preactivation.

## Methods

### Data and preprocessing

All analyses were conducted using freely available data from Demarchi et al.^3^ at this url: https://doi.org/10.5281/zenodo. It corresponds to a downsampled (to 100 Hz) and bandpassed filtered (1-30 Hz) version of the original MEG raw data which has been preprocessed using a signal space separation algorithm implemented in the Maxfilter program provided by the MEG manufacturer Elekta. The extensive methods about participants, experimental procedures, MEG data acquisition and preprocessing can be found in Demarchi et al.^3^

### Decoding and statistics

Data were analyzed with multivariate linear modeling algorithms available in MNE-python 1.5.1^9^ and scikit-learn 1.2.1^10^ (note that the original study used different software). We implemented several analytical pipelines. A multiclass Linear Discriminant Analysis (LDA) was used to classify the four sounds (i.e., labels, classes) from the MEG data (i.e., each datapoint had dimension equal to the number of MEG channels). Classifiers were always trained and tested at the level of individual participants. Separate classifiers were used at each time sample every 10ms.

Correlations between classification performance and sequence predictability were calculated for each participant independently, then Fisher-transformed. The Pearson coefficient was chosen instead of Spearman’s to remain as close as possible to the original linear regression approach. Time points at which classifiers performed significantly better than 0.25 (chance level chosen by Demarchi et al., corresponding to one divided by the number of classes), or at which correlations were significantly different from 0, were detected using non-parametric cluster-based permutation tests^11^ to adequately control for multiple comparisons while accounting for the statistical dependence between neighboring time points. We used the implementation from MNE-python^9^.

In addition, we ran Bayesian one-sample t-tests on the difference in Fisher-transformed correlation coefficients between the replication of the original study (pipeline 1 below) and our null approaches (pipelines 2 and 3). We used the R package BayesFactor (version 0.9.12-4.6) and its default prior on standardized effect sizes (Cauchy prior with scale = 2/2), designed to assign low prior probability to implausible effect sizes.

### Pipeline 1 (Figure 1c & 1e, Replication)

To reproduce the main results from Demarchi et al.^3^, the continuous data was segmented from 700 ms before to 700 ms after sound or omission onset. Omission trials and trials that were preceded by an omission trial were removed. In this first analysis, classifiers were trained only during the random block and tested during each of the four entropic conditions (random, midminus, midplus, ordered). A stratified 5-fold cross-validation scheme was used, meaning that at each cross-validation iteration, classifiers were trained on 80% of the random trials and separately tested on 20% of the trials in each block, preserving classes’ proportions. Using the same cross-validation scheme for each entropy (i.e., always testing on only 20% of the trials even if we could have used all trials when testing on an ordered sequence because training and testing trials are independent in this case) is better for comparing the decoding performance between entropies. A classifier that has been trained at a specific time sample was tested at all possible time samples using the temporal generalization method, resulting in two-dimensional arrays of decoding performance. The metric used to assess classifiers performance was the percentage of correct predictions. In a second analysis on omission trials, classifiers were also trained on all random sounds and tested only on omission trials in the four entropies. In all cases, training and testing partitions always contained different trials and did not overlap.

### Pipeline 2 (Figure 1c, Empirical Null)

To create a proper control condition where the null hypothesis of no anticipatory feature-specific activation would be true, continuous data of the random block were cropped and reordered to match the sequences in other entropy levels (midminus, midplus and ordered). In practice, to match the ordered sequence for example, random trials were cropped between 0 ms and 330 ms and reordered to match the ordered sequence of the same participant. We iterated across sound labels in the ordered sequence of the participant and selected the first occurrence of this sound in their random sequence and then removed this trial from the available random trials to be reordered. Such iteration always stopped before arriving at the last sound of the more ordered sequence. We used the same number of trials for all conditions, corresponding to the condition with the lowest number of reordered trials (2995 ± 59 trials on average). In order to be able to compare the results of the pipeline 1 (original ordering) and the pipeline 2 (reordering of the random data) (Figure 1), the same number of trials were used in both analytical pipelines. The same cross-validation and scoring steps from pipeline 1 were used in pipeline 2. Cross-validation folds were defined after reordering to ensure that the training and testing sets do not overlap (see script for details).

### Pipeline 3 (Figure 1e, Theoretical Null)

To reproduce the main results of Demarchi et al.^3^ using theoretical accuracy scores based on the transition matrices of each sequence and the confusion matrices of the classifiers, we extracted the confusion matrices at each training and testing times. Then, we used these confusion matrices to predict the behavior of the classifier in structured sequences under the null hypothesis of no predictive information in neural signals.

Let *C* be the confusion matrix of the classifier trained in the random sequence, i.e. *C*_*ij*_ is the probability that a true stimulus *i* will actually be decoded as stimulus *j* by the classifier.

Let *T* denote the matrix of transition probabilities between consecutive stimuli in a structured sequence. *T* encodes a simple Markov process, i.e. *T*_*ij*_ is the probability that stimulus *i* is followed by stimulus *j*:

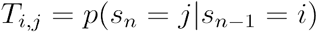

Note that, because all sounds are marginally equiprobable by design, the probability above can be rewritten using the Bayes law as:

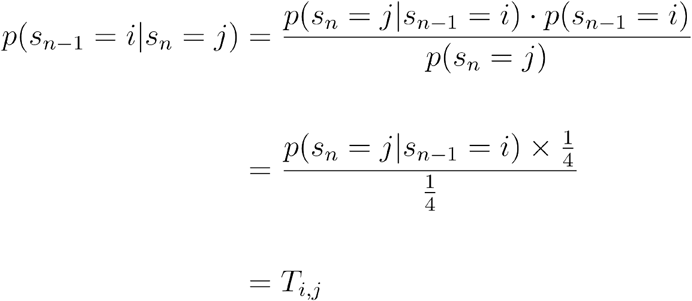

Now, if the target stimulus *n* is *j*, then the probability that it is decoded by chance (i.e. under the null hypothesis *H*_0_) from the preceding trial *n* ™ 1 can be calculated by marginalizing over the identity of the preceding sound. As a result, chance pre-stimulus decoding accuracy appears to depend both on the conditional probability distribution of the preceding stimulus described in the transition matrix, and on the associated terms in the confusion matrix:

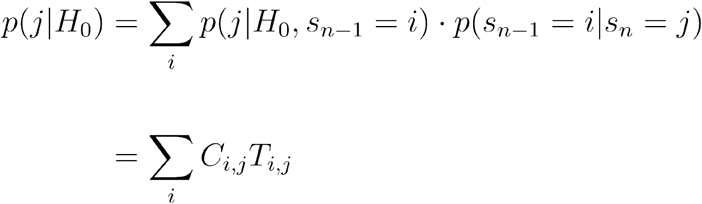

Integrating this term over all possible stimuli and taking into account that stimuli are equiprobable, the probability of the classifier to correctly predict the subsequent stimulus by chance only is:

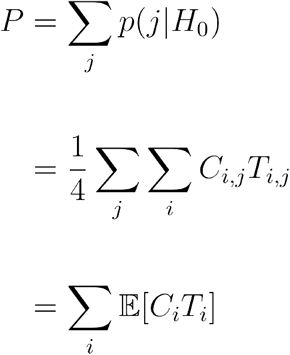

Therefore, for a structured sequence of stimuli, the chance level is calculated as the row-wise sum of the average products of corresponding terms from the transition and confusion matrices.

### Pipeline 4 (Figure 2)

In an attempt to increase the effect size of a putative feature-specific anticipatory signature, we decoded the most likely, rather than actual, upcoming stimulus. More specifically, we replicated pipelines 1 and 2, introducing a single change: the reference class used to determine whether the classifier is correct and calculate its accuracy was not the actual stimulus *n*, but the most probable identity of stimulus *n*, given the identity of *n* ™ 1 and the transition matrix ruling the sequence pattern (figure 1b).

## Code Availability

All MEG analysis pipelines in python, as well as the R code producing the statistics and the figures, are available at https://github.com/romquentin/predictive_activity/tree/main.

## Competing interests

The authors declare no competing interests.

## Author contributions

**Oussama Aboun:** Conceptualization, Formal analysis, Methodology, Visualization, Writing—original draft, Writing—review & editing

**Dmitrii Todorov:** Formal analysis, Writing—review & editing

**Arnaud Poublan-couzardot:** Formal analysis, Writing—review & editing

**Coumarane Tirou:** Writing—review & editing

**Antoine Lutz:** Writing—review & editing

**Marine Vernet:** Conceptualization, Writing—review & editing

**Romain Quentin:** Conceptualization, Formal analysis, Funding acquisition, Methodology, Project administration, Writing—original draft, Writing—review & editing

## Notes

### Competing Interest Statement

The authors have declared no competing interest.

### Funding

- Inserm: ATIP-Avenir granted to Romain Quentin
- UniversitM-CM-) Claude Bernard Lyon 1: ANR-11-LABX-0042

