## Supplementary Figures for "No evidence of neural feature-specific pre-activation during the prediction of an upcoming stimulus"

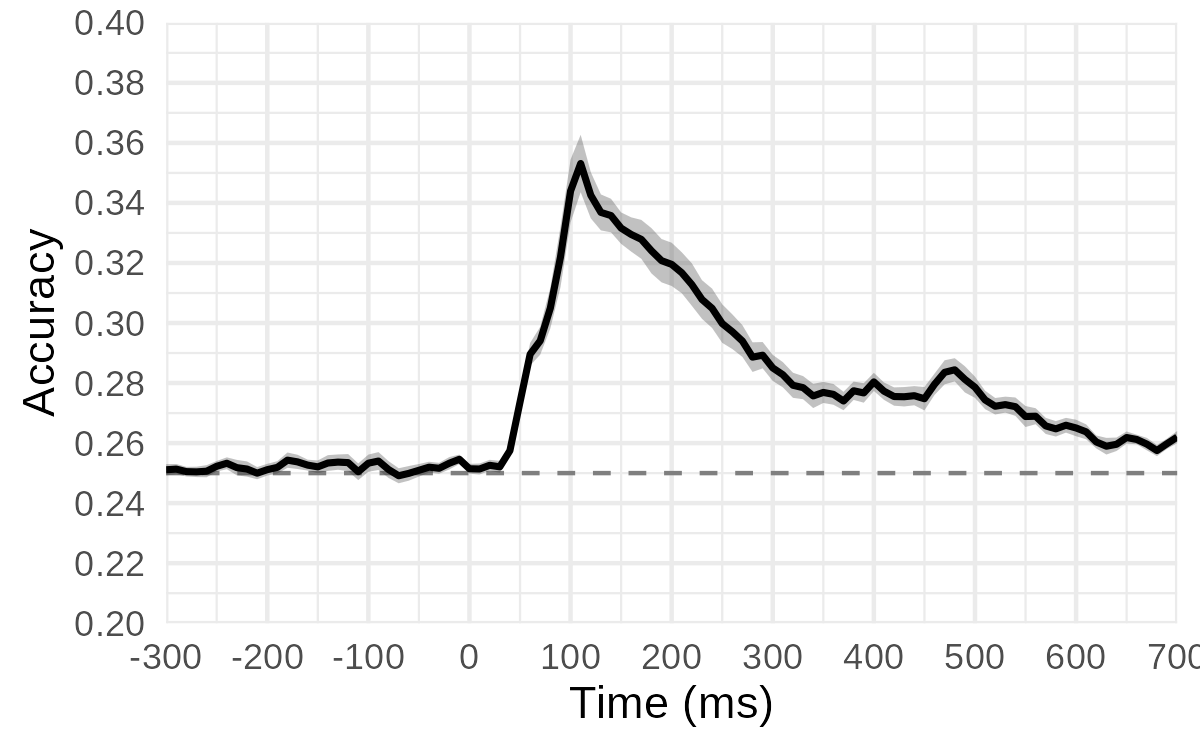

**Supplementary figure 1.** Decoding carrier frequencies from random sound sequences. This figure replicates figure 2a from the original publication.

**Supplementary figure 2.** Time-generalization decoding of omissions presented in structured sequences with a classifier trained on random sound sequences. **(a)** and **(b)** are analogous to figure 1c and 1d, respectively, but for omitted sounds.

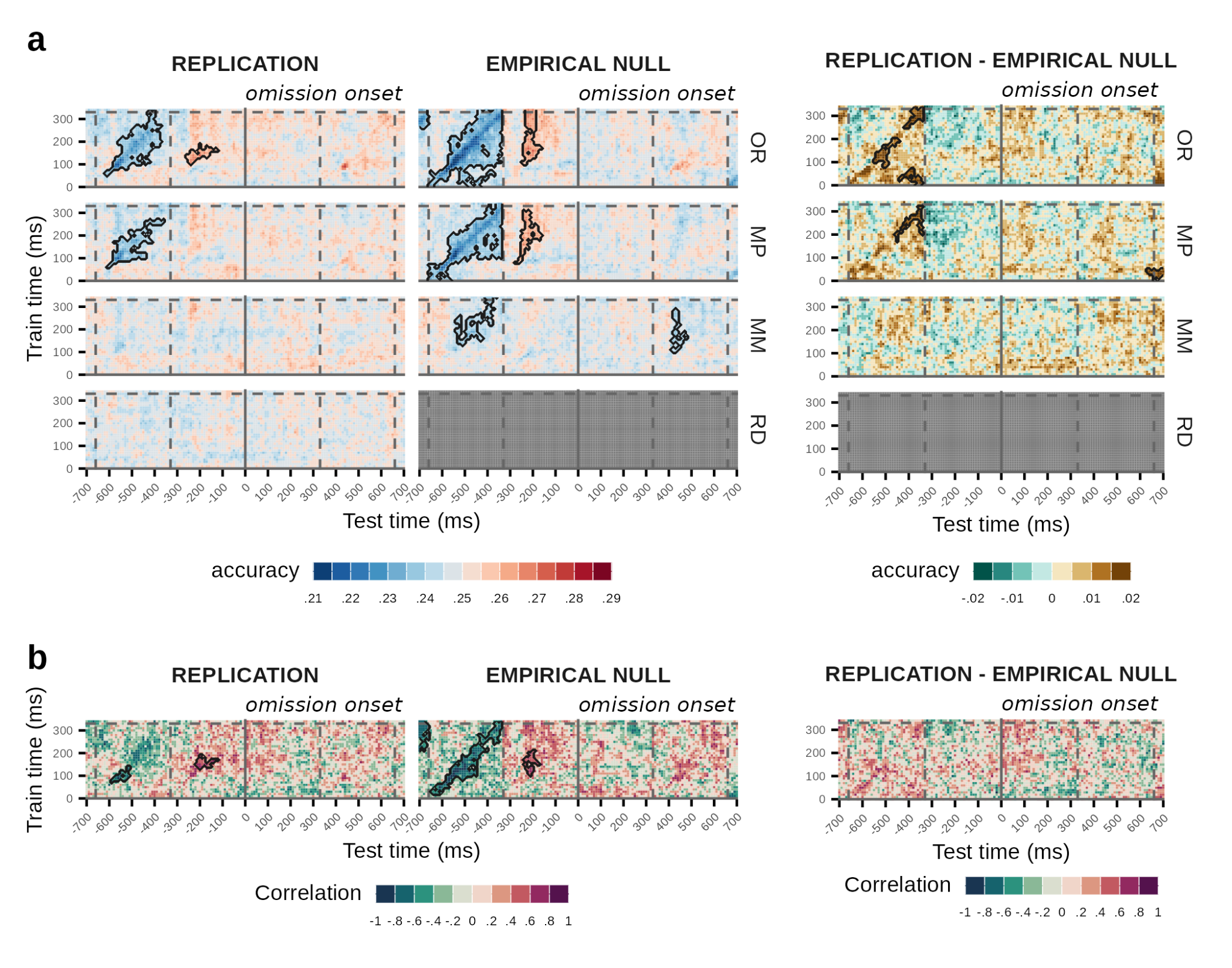

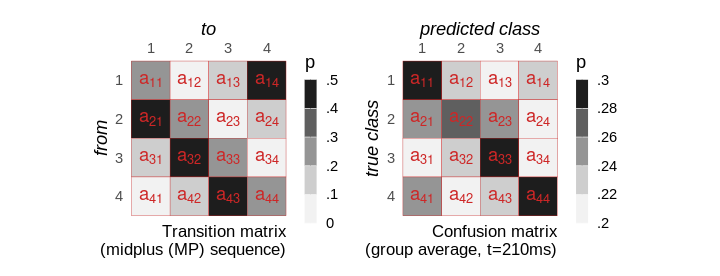

**Supplementary Figure 3.** The similarity between the transition and the confusion matrices induces spurious higher-than-chance decoding performance. **(a)** The descending nature of the pitch sequence manifests in elements of the first subdiagonal (a_21_, a_32_, a_43_) being higher than those of the second (a_31_, a_42_) and third (a_41_) sub-diagonals. The reverse pattern is found for super-diagonals. **(b)** Confusion matrices present partially similar structural features (here illustrated at 210 ms after stimulus onset): in particular, elements of the first subdiagonal are higher than those of the second one, and the fourth super-diagonal (a_14_) is higher than the third one (a_13_, a_24_). As a result, the two matrices show a weak but consistent positive correlation, inducing a decoding performance above 0.25 (as per equation 3) at trial $n-1$, even in the absence of anticipatory, feature-specific activation.

**Supplementary Figure 4.** Bayesian statistical analyses of the difference between true and synthetic data (top: empirical, bottom: theoretical), with respect to the accuracy-entropy correlation. Within most of the time window of the preceding stimulus (-333ms to 0ms), Bayesian Factors show moderate evidence in favor of the null hypothesis, including in the temporal cluster which was significant in the original data (delineated in black). Said otherwise, decoding results generated under the assumption of no preactivation of sensory signal are statistically similar to the original results.

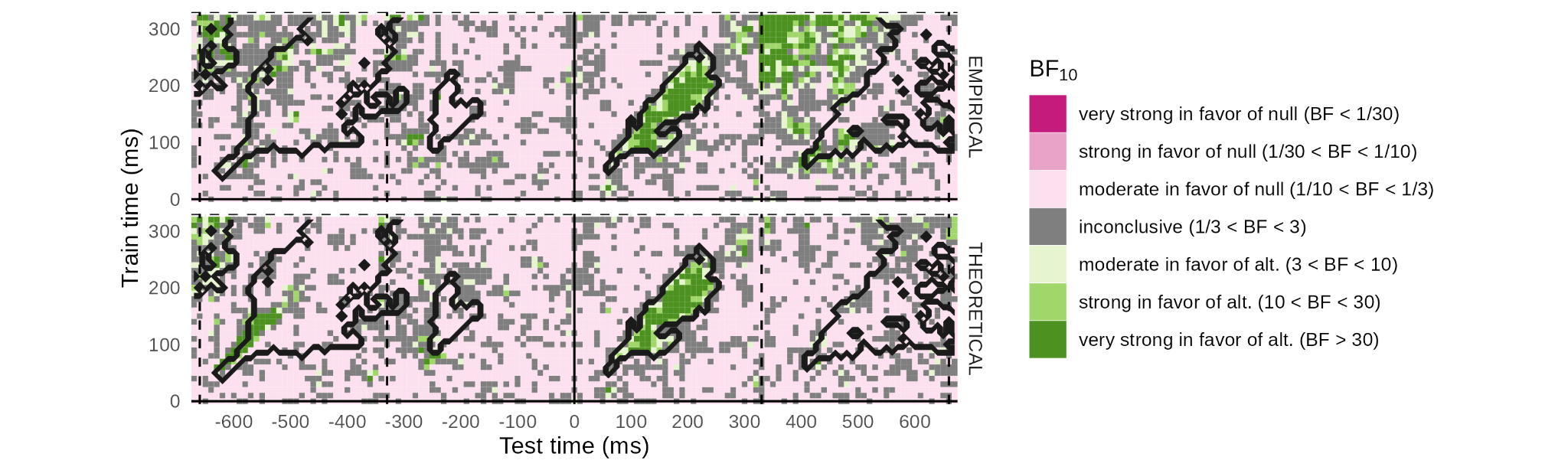
